# Decoding Surface Fingerprints for Protein-Ligand Interactions

**DOI:** 10.1101/2022.04.26.489341

**Authors:** Ilia Igashov, Arian R. Jamasb, Ahmed Sadek, Freyr Sverrisson, Arne Schneuing, Pietro Liò, Tom L. Blundell, Michael Bronstein, Bruno Correia

## Abstract

Small molecules have been the preferred modality for drug development and therapeutic interventions. This molecular format presents a number of advantages, e.g. long half-lives and cell permeability, making it possible to access a wide range of therapeutic targets. However, finding small molecules that engage “hard-to-drug” protein targets specifically and potently remains an arduous process, requiring experimental screening of extensive compound libraries to identify candidate leads. The search continues with further optimization of compound leads to meet the required potency and toxicity thresholds for clinical applications. Here, we propose a new computational workflow for high-throughput fragment-based screening and binding affinity prediction where we leverage the available protein-ligand complex structures using a state-of-the-art protein surface embedding framework (dMaSIF). We developed a tool capable of finding suitable ligands and fragments for a given protein pocket solely based on protein surface descriptors, that capture chemical and geometric features of the target pocket. The identified fragments can be further combined into novel ligands. Using the structural data, our ligand discovery pipeline learns the signatures of interactions between surface patches and small pharmacophores. On a query target pocket, the algorithm matches known target pockets and returns either potential ligands or identifies multiple ligand fragments in the binding site. Our binding affinity predictor is capable of predicting the affinity of a given protein-ligand pair, requiring only limited information about the ligand pose. This enables screening without the costly step of first docking candidate molecules. Our framework will facilitate the design of ligands based on the target’s surface information. It may significantly reduce the experimental screening load and ultimately reveal novel chemical compounds for targeting challenging proteins.

## 1 Introduction

Small molecule drug discovery is an arduous and resource-intensive process, characterised by declining productivity and high attrition rates. Computational design and, in particular, computational *structure-based drug design* (SBDD) — where the 3D structure of the target protein is known — of small molecules has been a successful strategy in developing novel therapies (Congreve et al., 2005). Typical computational SBDD workflows are based on *virtual screening* of large libraries of drug-like compounds, which are docked and scored by *scoring functions* to determine ideal geometries and complementarity (Ou-Yang et al., 2012). Various deep learning approaches have been applied to components of SBDD with promising results. However, rational *de novo* design has been relatively less studied by the machine learning community.

Fragment-based drug discovery (FBDD) has been a successful paradigm in early-stage drug development (Erlanson et al., 2016). The majority of the current workflows involve screening libraries of low molecular weight fragments against macromolecular targets of interest. Fragments can occupy multiple binding locations on the target providing starting points for developing candidate compounds through growing, linking or merging fragments. By incorporating structural knowledge of the target, a bottom-up approach like FBDD can result in reduced cost compared to high-throughput screening workflows (Erlanson et al., 2016). Furthermore, by designing candidate molecules around constituent favourable interactions, improved ligand efficiency can be achieved (Hopkins et al., 2014). However, fragments typically bind with much lower affinities than drug-like molecules making their identification and structural characterisation difficult.

Geometric deep learning, which encompasses deep neural networks operating on non-Euclidean domains (Bronstein et al., 2017), has shown considerable benefits in representation learning on proteins (Zhang et al., 2022; Jing et al., 2021; Gligorijević et al., 2021; Hermosilla et al., 2021) and other molecular structures (Fang et al., 2022; Townshend et al., 2021), particularly in the context of drug discovery (Gaudelet et al., 2021; Méndez-Lucio et al., 2021).

In this work, we develop a novel ligand and fragment searching methodology based on geometric deep learning applied to surface representations of binding pockets. We demonstrate the effectiveness of our method on a subset of the PDBbind dataset (Wang et al., 2004) and show that the proposed approach performs on par with the state of the art and outperforms a recent deep learning-based method. We also develop a binding affinity prediction model competitive with the current state of the art. Our method requires limited knowledge of the binding pose, accelerating virtual screening workflows by circumventing docking and screening of compounds and targets for which co-crystallised structures are not available.

### 1.1 Related Work

#### Pocket-Centered Ligand Screenin

Binding pocket similarity plays an important role in drug discovery. Incorporating structural knowledge about pockets of interest enables efficient search of relevant binding molecules and provides new means for exploring molecular space. Provided that a ligand can be recognized by different residues with different interaction types (Barelier et al., 2015), finding a suitable pocket representation is the most crucial and highly challenging part of the task. Various pocket representations have been explored, including pharmacophore (Wood et al., 2012) and deep learning-based fingerprints (Simonovsky & Meyers, 2020) as well as geometric and chemophysical descriptors (Konc & Janežič, 2010; Shulman-Peleg et al., 2004; Zhu et al., 2015). Recently, pocket-centered methods for conditional molecule generation (Masuda et al., 2020; Méndez-Lucio et al., 2021) have been proposed. Stärk et al. (2022) developed a geometric deep learning method for predicting the receptor binding location and the ligand’s bound pose and orientation.

#### Fragment-Based Drug Discover

In a standard setting, screening methods are fundamentally limited by the diversity of the underlying compound libraries. FBDD operates on small labile molecules (typically with molecular weight *<* 300 Da) to identify low potency, high-quality leads, which are then matured into more potent, drug-like compounds (Imrie et al., 2020). Millions of compounds such as rings, linkers and scaffolds are available in databases such as ZINC (Sterling & Irwin, 2015). However, identifying and filtering relevant fragments from such a database is a challenging problem (Kirsch et al., 2019). Instead, one can split molecules of interest into smaller compounds based on various chemical and geometric assumptions (Bemis & Murcko, 1996; Lewell et al., 1998; Degen et al., 2008; Ghersi & Singh, 2014; Jin et al., 2018) to generate a relevant fragment database. Another FBDD-specific problem is constructing a ligand from the found fragments. Various linking (Böhm, 1992; Thompson et al., 2008; Yang et al., 2020) and scaffold hopping (Maass et al., 2007; Vainio et al., 2013) methods have been developed to address this task. Recently, deep generative models have been proposed (Jin et al., 2018) or adopted (Liu et al., 2018; Imrie et al., 2020) for these purposes. Notably, Drotár et al. (2021) developed a structure-based method for generating drug-like ligands from molecular fragments via a sequential VAE-based approach, conditioning each step of the generative process on a graph representation of the binding pocket.

#### Binding Affinity Prediction

Predicting the affinity of a given protein-ligand interaction is an important component of virtual screening pipelines. Traditionally, methods for predicting binding affinity have been based on physics-based free energy calculations which do not account for entropic contributions (Kitchen et al., 2004). Machine learning approaches based on hand-crafted features, which do not impose a pre-defined functional form, were shown to improve performance by learning from the experimental data (Ballester & Mitchell, 2010). Several types of deep learning models have been applied to this problem, including 3D-CNNs (Stepniewska-Dziubinska et al., 2018; Jiménez et al., 2018; Zheng et al., 2019), GNNs (Li et al., 2021), and surface-based models (Somnath et al., 2021). However, the majority of existing ML-based approaches heavily rely on co-crystallised structures of complexes which are hard and not always possible to obtain. Often, docking algorithms are used which typically result in noisier and more error-prone poses that prove more challenging to accurately predict binding affinity from (Jones et al., 2021). Furthermore, these approaches are insensitive to scenarios where multiple binding mechanisms and poses contribute to the overall affinity (Stjernschantz & Oostenbrink, 2010).

#### Protein Surface Representation

Structure-based encoders of protein molecular surfaces have been successfully applied to predicting protein-protein interaction sites and identifying potential binding partners in protein docking (Gainza et al., 2019; Sverrisson et al., 2021; Somnath et al., 2021). Molecular surface interaction fingerprinting (MaSIF) (Gainza et al., 2019) pioneered learning geometric surface descriptors for solving protein interaction-related tasks. It computes geometric and chemical features on a triangulated surface mesh and aggregates local information using a geometric deep learning architecture based on geodesic convolution operators (Bronstein et al., 2017). Differential MaSIF (dMaSIF) (Sverrisson et al., 2021) is a derived method that sidesteps the expensive mesh generation and feature pre-computation steps and creates a lightweight point cloud representation of the molecular surface solely from raw atom coordinates and types. Feature vectors at each surface point are updated by applying approximate geodesic convolutions resulting in the final *embedding* vectors that can be further used in various downstream tasks. In this work, we employ these vectors to compare protein binding pockets and to predict protein-ligand binding affinity. Figure 1 illustrates the workflow of dMaSIF.

**Figure 1:**
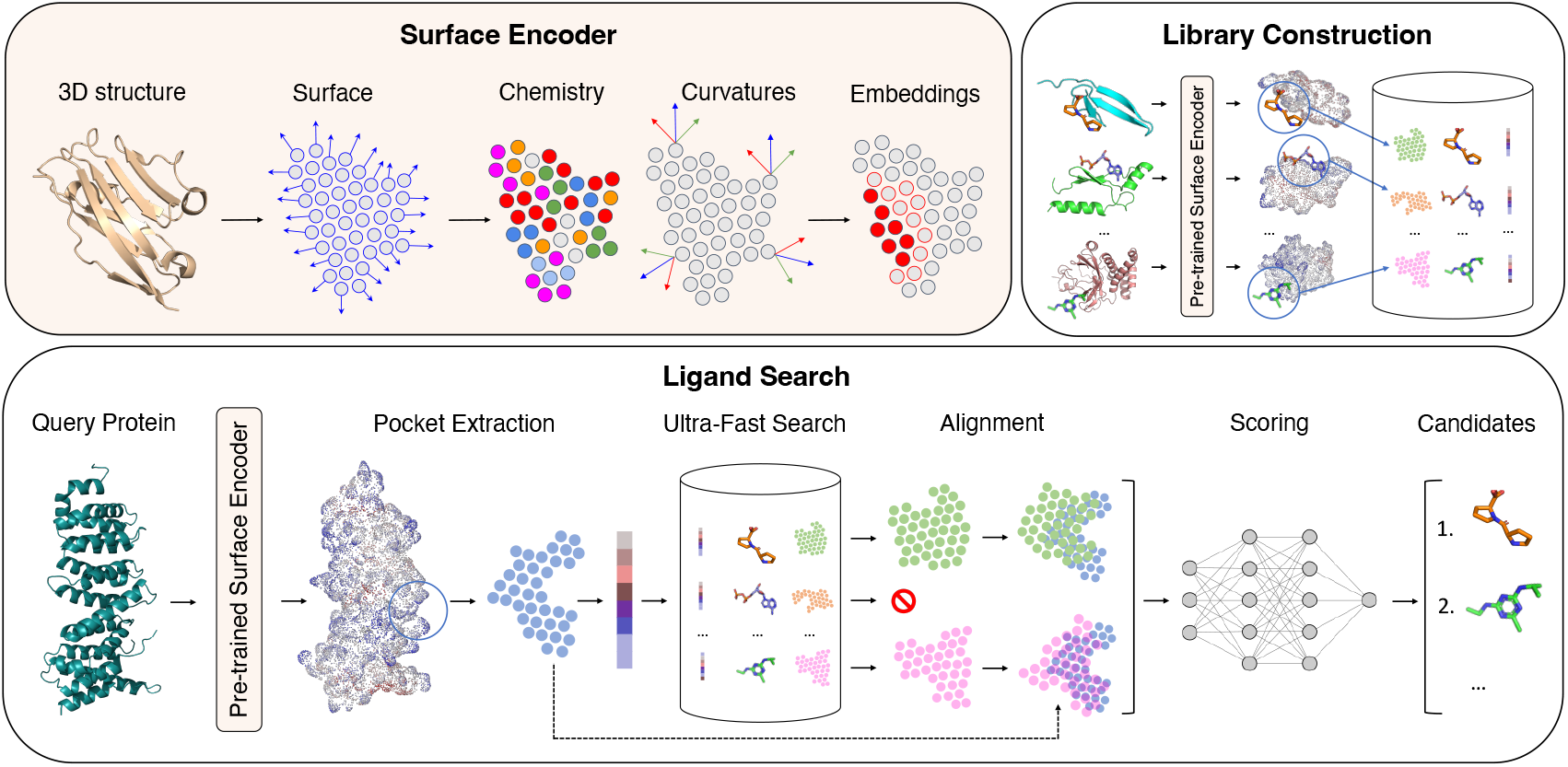
Surface encoding workflow of dMaSIF (top left), library construction (top right), and ligand search pipeline (bottom).

## 2 Methods

### 2.1 Ligand Search

A typical pocket-centered drug discovery workflow involves screening libraries containing millions of molecules to identify an initial set of candidates. To ensure effectiveness, both a suitable pocket description and appropriate search procedure are required. In this section, we introduce a novel pocket-centered ligand search pipeline. We construct a *library* of available (known) ligand-pocket pairs that are queried by new (*query*) pockets to identify candidate ligands that are most likely to bind to the input pockets.

#### Dataset Preparation

We consider a subset of PDBbind v2020 (Wang et al., 2004) by combining the general (*n* = 14, 127) and refined (*n* = 5, 316) sets and selecting only complexes with ligands that are not amino acids, which are subsequently parsed by RDKit (Landrum et al., 2013), have QED *>* 0.3 (Bickerton et al., 2012), and are present in more than 10 complexes. We obtain 488 complexes and protonate the structures with Reduce (Word et al., 1999). To construct the query (*n* = 130) and library (*n* = 358) datasets, we compute sequence similarity clusters for all proteins using CD-HIT (Fu et al., 2012) with a similarity threshold of 0.9. For each ligand, we split the set of the corresponding complexes in query and library sets in proportion 30*/*70, such that clusters assigned to each set do not overlap.

#### Protein Pockets

To retrieve the protein pocket, we first generate a surface of the protein and compute embeddings for each point on the surface using dMaSIF (Sverrisson et al., 2021). For each ligand atom, we select the closest point on the protein surface. For each selected point, we consider its *r-neighborhood*, a set of points within Euclidean distance *r*. We consider different values for *r* (see Appendix A), and achieve the best performance with *r* = 2 Å. The resulting pocket embedding is generated as a union of *r*-neighborhoods of all selected points.

#### Workflow

For each pocket in the query set, we *search* for the most similar binding pockets in the library and output the corresponding ligands as candidates for the query pocket. The general scheme of the workflow is represented in Figure 1 and consists of three main steps: shortlisting, alignment and scoring. First, we perform *ultra-fast search* over the whole library in order to shortlist candidate ligands. The purpose of this step is to filter out irrelevant ligands and hence reduce the computational complexity of the subsequent steps. For each query pocket, we select the top-50 candidates based on *similarity* between *global* pocket embeddings. As shown in Appendix A, we consider two similarity functions: Euclidean distance and dot product. We achieve the best performance using Euclidean distance. Global pocket embeddings are created by averaging the embedding vectors of all points in the pocket. An alternative approach was to consider an embedding vector of a single point, e.g. of the one closest to the center of the pocket. However, as shown in Appendix A, better results were achieved with averaging. In the second step, we align pockets of shortlisted candidates with the query pocket. We consider two methods: Random Sample Consensus (RANSAC) (Fischler & Bolles, 1981) followed by point-to-point Iterative Closest Point (ICP) (Besl & McKay, 1992), and our own optimization-based alignment approach described in Section 2.2. Once the shortlisted pockets are aligned with the query pocket, we score pairs of pockets using a pre-trained neural network (described in Section 2.3).

### 2.2 Optimization-Based Alignment

Consider two point clouds: a source point cloud with *M* points described by coordinates 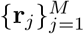, normal vectors 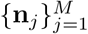, and embedding vectors 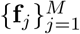, and a target point cloud with *N* points described by coordinates 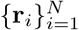, normal vectors 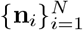, and embedding vectors 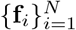.

We represent normal vectors and embeddings of the target point cloud as continuous vector fields 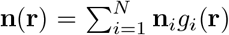 and 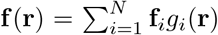 correspondingly in ℝ^3^ by smoothing the discrete points of the target point cloud via Gaussian kernels:

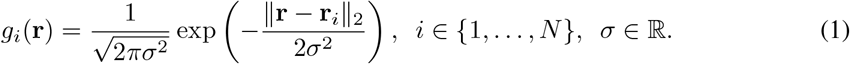

Our task is to find the optimal orientation of the source point cloud with respect to the vector fields **f** (**r**) and **n**(**r**). The best orientation of the source point cloud is the orientation that maximizes two terms. The first term is similarity *S* between embedding vectors of the source point cloud and values of the field **f** (**r**) in coordinates that correspond to the positions of points of the source point cloud. Similarity can be expressed as Euclidean distance, dot product, or cosine between two vectors. The second term is dot product between normal vectors of the source point cloud and values of the field **n**(**r**) in coordinates that correspond to the positions of points of the source point cloud. Orientation of the source point cloud is parameterized by rotation matrix **R** ∈ *M*_3_(ℝ), **R**^⊤^**R** = 𝕀, det **R** = 1, and translation vector **t** ∈ ℝ^3^. The objective function can be written as follows,

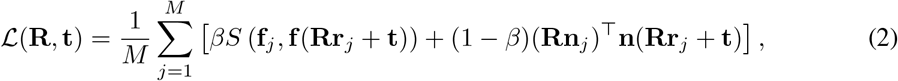

where *β* ∈ [0, 1] is a parameter that controls the contribution of each term in the final objective value. Using a 6D-parameterization **R** = **R**(**u, v**) of rotation matrix **R** with **u** ∈ ℝ^3^, **v** ∈ ℝ^3^ as described in Zhou et al. (2019), the optimization problem can be written as max 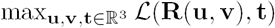.

To find the optimal rotation and translation parameters, we perform gradient descent in ℝ^3^ ×ℝ^3^× ℝ^3^ implemented in PyTorch (Paszke et al., 2017) using Adam optimizer (Kingma & Ba, 2014).

### 2.3 Scoring Neural Network

For the final ranking after pocket alignment, we train a neural network to distinguish between good — where pockets correspond to the same ligand and are properly aligned — and bad matches. The training dataset is crucial to the method’s success as it implicitly defines a notion of quality that is subsequently used to score shortlisted pairs of pockets.

#### Dataset

We prepare data in a similar procedure to Section 2.1. However, here we select ligands represented in fewer than ten complexes. This means that complexes used for training are not used in the ligand search experiments. To create positive training examples we select pairs of pockets that bind to the same ligands but originate from complexes with less than 90% sequence identity. We align these pockets based on ligand atom coordinates using rigid-body SVD-based alignment (Arun et al., 1987). To create negative examples we randomly select pairs of pockets that correspond to different ligands and match their centers of mass. We randomly split the positive and negative example pairs into training (*n* = 2, 562) and validation (*n* = 654) sets.

#### Input Features

For a given pair of pockets, we first compute their point-wise features that are input to the network. For each point in the source pocket we find the closest point in the target pocket, and for each of the resulting point pairs, we compute three values: inverted distance between two points, dot product of normal vectors, and dot product of embedding vectors.

#### Architecture

The scoring model consists of two symmetric blocks with global average pooling between them. The first block takes three-dimensional feature vectors for each point and projects them to the 256-dimensional hidden vector via three fully-connected layers followed by batch normalization (Ioffe & Szegedy, 2015) layers and ReLU activations. Global pooling averages hidden vectors over all points in the source pocket resulting in a single 256-dimensional vector which is processed by the final block. This block includes a sequence of three fully connected layers with ReLU activations followed by softmax. For symmetry, each pocket pair is processed twice: with the original order of source-target pockets and with the swapped order. Final predictions are averaged.

### 2.4 Fragment Search

Fragment-based search has a notable advantage compared to the ligand search. Although the number of unique available ligands is usually orders of magnitude larger than the number of unique constituent fragments, the latter can be considered as building blocks of ligands. Therefore, the fragment-based method becomes less dependent on the available data and allows exploration of ligand space by combining available fragments. Below, we describe the modified search approach that operates on fragments instead of entire ligands.

#### Fragment Generation

To decompose ligands into fragments, we use BRICS (Degen et al., 2008) which simultaneously breaks retrosynthetically relevant bonds and filters unwanted chemical motifs and small terminal fragments. For each ligand in the dataset described in Section 2.1, we retrieve its BRICS fragments, remove open exits, and filter out fragments with molecular mass *>* 300 Da, deduplicate the resulting set based on Tanimoto distance, and match every resulting compound with other ligands in order to get all its occurrences. In total, we gather 36 unique fragments. Exemplary fragments along with their occurrences are provided in Figure S4. Mapping unique fragments back to ligands in which they occur, we obtain the query (440 entries) and library (1,434 entries) datasets for fragment search.

#### Workflow

The fragment search process is very similar to the ligand-based algorithm described in Section 2.1 and fragment pockets are constructed in the same way. However, as fragments are smaller, the resulting point clouds constructed for fragments describe smaller regions of ligand binding pockets. For clarity, we refer to these regions as fragment *patches*. The main difference between ligand and fragment search workflows lies in the first step, ultra-fast search. In ligand search, the top-ranked candidates in the library are returned. In fragment search, for each fragment in the library (that can have more than one corresponding patch), we select a single patch that has the highest similarity score among all patches corresponding to a given fragment. This allows us to increase the diversity of candidates in the subsequent steps of the pipeline.

### 2.5 Binding Affinity Prediction

#### Dataset Preparation

We use PDBbind v2016 (Liu et al., 2017), a dataset of protein structures co-crystallised with ligands and their associated binding affinity. Structures are protonated with Reduce (Word et al., 1999) and only atoms belonging to the polypeptide chain(s) are extracted. To enable comparison with existing methods, the training set is constructed from the refined set, from which we randomly sample 10% for the validation set. Models are evaluated on the CASF 2016 core set (Su et al., 2018) (which is removed from the training data). Binding pocket embeddings are extracted as described in Section 2.1 using a much larger neighbourhood radius of *r* = 12 Å to reduce the sensitivity to pose. Ligands are represented as molecular graphs. Edges correspond to chemical bonds (single, double, triple, aromatic or ring) and node features include one-hot encoded atom types, degree, valence, hybridization state, number of radical electrons, formal charge, and aromaticity. Crucially, we make no use of positional information in the ligand representation.

#### Architecture

We train a model using a dMaSIF-based encoder for the protein pocket. Importantly, the encoder receives the whole protein surface. After encoding, pocket point embeddings are extracted and aggregated into a global embedding vector by taking the element-wise max of individual point embeddings as we found this resulted in better performance than sum or average pooling. As we focus on understanding the effectiveness of surface-based descriptors, we use a simple GCN (Kipf & Welling, 2016) as the ligand encoder and aggregate node embeddings via sum pooling to obtain the graph representation. Ligand and pocket embeddings are concatenated and used as input for an MLP decoder to predict the binding affinity values. Hyperparameters are provided in Table S11.

## 3 Results

### 3.1 Ligand Search

In this experiment, we evaluated the ability of the proposed multi-staged search algorithm to output relevant ligands to the query pockets. We compare our approach with three state-of-the-art methods for pocket-centered ligand screening, ProBiS (Konc & Janežič, 2010), KRIPO (Wood et al., 2012), and DeeplyTough (Simonovsky & Meyers, 2020). ProBiS detects structurally similar sites on protein surfaces by local surface structure alignment using an efficient maximum clique algorithm (Konc & Janežič, 2010). KRIPO is a method for quantifying the similarities of binding site subpockets based on pharmacophore fingerprints (Wood et al., 2012). DeeplyTough is a convolutional neural network that encodes a three-dimensional representation of protein pockets into descriptor vectors that are further compared using Euclidean distance (Simonovsky & Meyers, 2020). We note that KRIPO failed on 6 query pockets, hence all the resuts discussed below were obtained on 124 query pockets successfully processed by all the methods.

In Table 1, we report the fractions of pockets from the query set for which the methods returned correct ligands in top-1, top-5, top-10, top-20, and top-50. Our best performing method operates on pockets built with neighborhood radius *r* = 2 Å and using a dMaSIF-search model trained (for 33 epochs) with the standard parameters (Sverrisson et al., 2021) except subsampling (set to 150), resolution (set to 0.7 Å), and embedding size (set to 16). We report metrics computed after the first and last stages of the search process. Namely, metrics were computed for lists of candidates ranked by global search score and neural network score. We provide results of pipelines that use RANSAC and our optimization method for alignment.

**Table 1:** Ligand search results.

| Method | Top-1 | Top-5 | Top-10 | Top-20 | Top-50 |
| --- | --- | --- | --- | --- | --- |
| ProBiS (Konc & Janežič, 2010) | **0.581** | 0.718 | 0.750 | 0.806 | 0.863 |
| KRIPO (Wood et al., 2012) | 0.573 | **0.726** | **0.798** | **0.863** | **0.935** |
| DeeplyTough (Simonovsky & Meyers, 2020) | 0.331 | 0.597 | 0.710 | 0.758 | 0.903 |
| Ours: search | 0.387 | 0.516 | 0.629 | 0.734 | **0.871** |
| Ours: search+ransac+score | **0.516** | **0.661** | **0.742** | **0.823** | **0.871** |
| Ours: search+optim+score | **0.516** | 0.637 | 0.710 | 0.766 | **0.871** |

Our full-scale search pipeline performs on par with state-of-the-art methods (Table 1). Notably, the first stage of the search process, ultra-fast global search, returns relevant ligands within the top-50 candidates in 87% of cases, making it appropriate for the initial shortlisting of candidates prior to fine-grained scoring.

#### Pocket Clustering

To illustrate that global embeddings of pockets contain information about the types of binding ligands, we select pockets bound by five structurally different ligands, compute all-by-all pairwise similarities between pairs of their cognate binding pockets, and visualize the results using multidimensional scaling (Torgerson, 1952). As pockets corresponding to the same ligand should show functional and structural similarity, we expect the embeddings of these pockets to be clustered in the plane. The embeddings of the selected pockets should be grouped in 5 clusters, where each cluster corresponds to a certain ligand. The resulting distribution of points is shown in Figure 2A and clearly demonstrates that pockets binding to the same ligands are grouped together.

**Figure 2:**
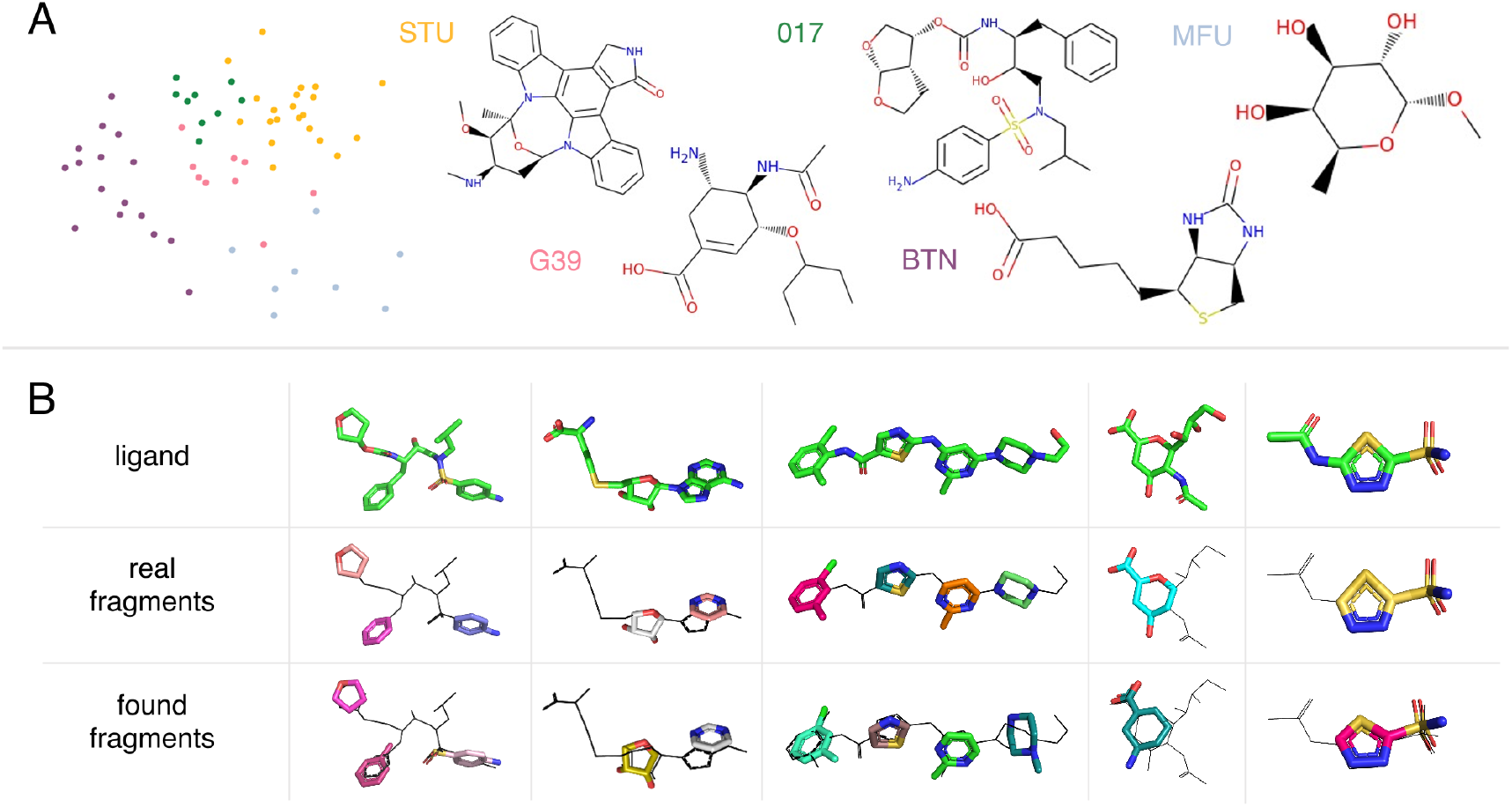
(A) Multidimensional scaling applied to pairwise similarities of pockets corresponding to five different ligands. Each point represents a single pocket coloured by the corresponding ligand type. (B) Fragment search examples. For each ligand, we show the most suitable found and aligned fragments from top-10. Black lines represent parts of ligands that are not covered by fragments.

### 3.2 Fragment Search

We evaluate our fragment search algorithm in the same way as our ligand-search approach (Section 3.1). Instead of ligand-pocket pairs, we consider fragment-patch pairs, and search for candidates for each fragment’s patch separately. The scoring neural network explained in Section 2.3 was retrained specifically for scoring patches of fragments. We compute Pearson and Spearman correlation of the predicted scores with Tanimoto similarity scores between fragments, and compare our method with KRIPO. We note that KRIPO failed on 16 query patches, hence the results discussed below were obtained on 424 query patches successfully processed by all the methods. Table 2 summarizes the performance of the fragment search pipeline. It includes results obtained after the ultra-fast search stage and after the scoring step that processed patches aligned with RANSAC and our optimization-based algorithm. Figure 2B contains several ligands with their ground-truth fragments and fragments identified and aligned with RANSAC. In each case, fragments were manually selected from the top-10 candidates returned by the algorithm.

**Table 2:** Fragment search results.

| Method | Top-1 ↑ | Top-5 ↑ | Top-10 | Top-20 | Pearson | Spearman |
| --- | --- | --- | --- | --- | --- | --- |
| KRIPO (Wood et al., 2012) | **0.469** | **0.724** | 0.816 | 0.873 | 0.089 | 0.098 |
| ours:search | 0.255 | 0.717 | **0.892** | **0.958** | **0.346** | 0.127 |
| ours:search+ransac+scoring | 0.245 | 0.594 | 0.715 | 0.861 | 0.119 | 0.126 |
| ours:search+optim+scoring | 0.245 | 0.479 | 0.601 | 0.776 | 0.172 | **0.136** |

### 3.3 Affinity Prediction

To assess the ability of the model to learn useful representations of protein pocket surfaces in the context of small-molecule binding, we train a binding affinity predictor. We are concerned with the scenario in which a co-crystallised structure is unavailable, as this is the most likely scenario in practice. We therefore compare our approach to baseline methods evaluated on docked poses of ligands in the PDBbind v2016 core set and demonstrate state-of-the-art performance without requiring accurate docking as an initial step (Table 3). We also compare to baseline models evaulated on poses from co-crystallised structures and perform on par with the alternative methods despite not making use of pose information (Table S12).

**Table 3:** Binding affinity prediction performance on PDBbind v2016 Core Set Docking Poses. [*] results taken from Jones et al. (2021).

| Method | RMSE ↓ | MAE ↓ | R ↑ |
| --- | --- | --- | --- |
| SG-CNN* | 1.576 | 1.277 | 0.699 |
| 3D-CNN* | 2.558 | 2.058 | 0.537 |
| Vina* (Trott & Olson, 2009) | - | - | 0.616 |
| MM-GBSA* | - | - | 0.629 |
| FAST-midlevel* (Jones et al., 2021) | 1.874 | 1.487 | 0.702 |
| FAST-late* (Jones et al., 2021) | 1.871 | 1.498 | 0.712 |
| Ours | **1.540** | **1.248** | **0.721** |

## 4 Conclusion

Accurate prediction of small-molecule protein interactions remains a very challenging task for computational methods. Here, we propose a general framework that leverages protein surface descriptors for small molecule related tasks. Our novel ligand and fragment searching methods can be employed as starting points in FBDD or as initialisations to generative chemistry models to develop novel chemical matter in a principled structure-based manner. Furthermore, we underline that this approach has been developed using a surface embedding model trained for predicting protein-protein interactions. There is significant scope to improve performance by developing surface encoders explicitly trained on tasks more closely related to protein-ligand interactions. We emphasize that properties of the surface embedding space play a crucial role in the ability to identify similar pockets based on dot product or distance similarity metrics. Therefore, an ideal protein surface encoder for this task should be trained in a way that it constrains the resulting embedding space to be Euclidean. Additionally, we develop a binding affinity predictor that is comparable in performance to existing methods without explicit consideration of pose or modelling of intramolecular interactions. We note that incorporating pose information is a natural extension of our work though this framing does not retain some of the reduced pose sensitivity advantages we sought to. The components discussed in this paper can be combined in subsequent work to use a fully-differentiable affinity predictor as a scoring function to directly optimize fragment placement. The resulting set of fragments can be further merged into a single chemically relevant molecule.

## Acknowledgments

This project has received funding from the European Union’s Horizon 2020 research and innovation programme under the Marie Skłodowska-Curie grant agreement No 945363. ARJ is funded by the Biotechnology and Biological Sciences Research Council (BBSRC) DTP studentship (BB/M011194/1). TLB. thanks the Wellcome Trust for support through an Investigator Award (200814/Z/16/Z; 2016-2021). MB is supported in part by the ERC Consolidator Grant No. 724228 (LEMAN).

We thank Charlie Harris, Casper Goverde, Leo Scheller, and Andreas Scheck for helpful discussions.

## A Ligand Search Parameters

### Pocket construction

The radius of the neighborhood used for extracting binding pockets plays an important role in the performance of the global search. Fixing the remaining parameters, we experiment with different neighborhood radii (shown in Table S4) and identify an optimal radius of *r* = 2 Å.

**Table S4:**
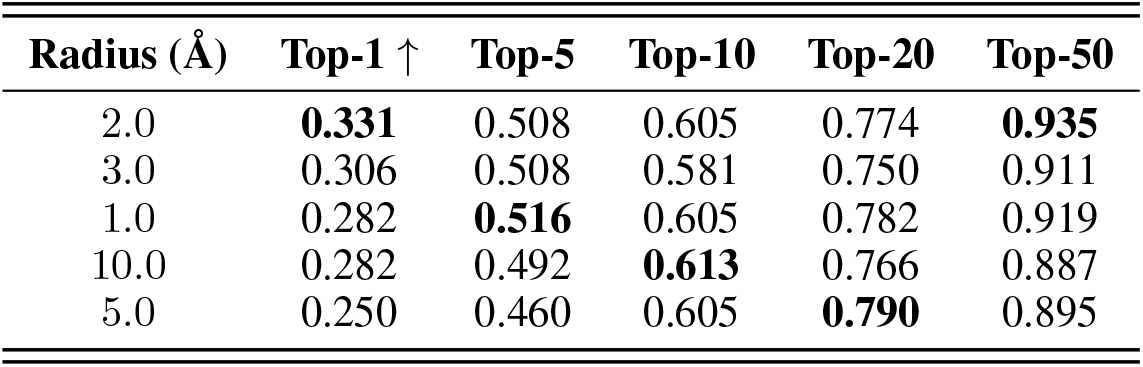
Global ligand search experiments with different neighborhood radius values.

### Global search

To find the best way of constructing global pocket embeddings, we experimented with two aggregation schemes: simple averaging and taking the embedding vector of the point closest to the center of mass of the pocket. For similarity measures, we consider Euclidean distance and dot product. Fixing the remaining parameters, we perform global search experiments with different aggregation and similarity functions (Table S5) identifying mean aggregation and Euclidean distance similarity as the most performant schemes.

**Table S5:** Global ligand search experiments with different aggregation and similarity functions.

| Averaging | Similarity | Top-1 ↑ | Top-5 | Top-10 | Top-20 | Top-50 |
| --- | --- | --- | --- | --- | --- | --- |
| Mean | Euclidean | **0.331** | 0.508 | 0.605 | 0.774 | **0.935** |
| Center | Euclidean | 0.274 | **0.548** | **0.685** | **0.815** | 0.887 |
| Mean | Dot | 0.024 | 0.024 | 0.113 | 0.274 | 0.452 |
| Center | Dot | 0.024 | 0.194 | 0.306 | 0.444 | 0.613 |

### dMaSIF

The choice of the pre-trained dMaSIF model plays an important role. We consider dMaSIF models pre-trained for three different tasks: protein-protein interaction search (dMaSIF-search), protein-ligand binding affinity prediction (dMaSIF-affinity), and protein-ligand pocket classification (dMaSIF-ligand). An important dMaSIF property that we assume should matter in our task is granularity of surfaces produced by dMaSIF. This property is controlled by two hyperparameters: *subsampling* and *resolution*. Subsampling determines the initial number of points sampled around each atom during the first step of the dMaSIF pipeline (Sverrisson et al., 2021). Resolution controls the size of a 3D voxel that should contain not more than one point (Sverrisson et al., 2021). The lower the resolution, the more detailed the resulting surface will be. We consider two dMaSIF-search models: with subsampling 100 and resolution 1 Å, and with subsampling 150 and resolution 0.7 Å. Due to the high computational complexity, we had to set subsampling 20 and resolution 1 Å for dMaSIF-affinity. For the same reason, the number of training epochs differs for each dMaSIF model.

**Table S6:** Global ligand search experiments with different pre-trained dMaSIF models.

| Model | Subsampling | Resolution ($\text{\AA}$) | Top-1 ↑ | Top-5 | Top-10 | Top-20 | Top-50 |
| --- | --- | --- | --- | --- | --- | --- | --- |
| dMaSIF-search | 150 | 0.7 | **0.387** | **0.516** | 0.629 | 0.734 | 0.871 |
| dMaSIF-search | 100 | 1.0 | 0.331 | 0.508 | 0.605 | 0.774 | **0.935** |
| dMaSIF-affinity | 20 | 1.0 | 0.266 | **0.516** | **0.653** | **0.782** | 0.871 |
| dMaSIF-ligand | 100 | 1.0 | 0.242 | 0.395 | 0.516 | 0.661 | 0.823 |

Parameters for dMaSIF models listed in Table S6 are provided in Table S7. For training the models, we used the code from https://github.com/FreyrS/dMaSIF. In case of dMaSIF-ligand and dMaSIF-affinity, we slightly adjusted the code for new purposes. dMaSIF-search models were trained on the same dataset as in Sverrisson et al. (2021). dMaSIF-ligand was trained on the dataset that was used for training MaSIF-ligand Gainza et al. (2019). Dataset for dMaSIF-affinity is described in Section 2.5.

**Table S7:** Parameters of dMaSIF models used in this work.

| model | dMaSIF-search | dMaSIF-search | dMaSIF-ligand | dMaSIF-affinity |
| --- | --- | --- | --- | --- |
| embedding_layer | dMaSIF | dMaSIF | dMaSIF | dMaSIF |
| resolution | 0.7 | 1.0 | 1.0 | 1.0 |
| distance | 1.05 | 1.05 | 1.05 | 1.05 |
| variance | 0.1 | 0.1 | 0.1 | 0.1 |
| sup_sampling | 150 | 100 | 100 | 10 |
| atom_dims | 6 | 6 | 6 | 6 |
| emb_dims | 16 | 16 | 16 | 32 |
| in_channels | 16 | 16 | 16 | 16 |
| orientation_units | 16 | 16 | 16 | 16 |
| post_units | 8 | 8 | 8 | 8 |
| n_layers | 3 | 3 | 3 | 1 |
| radius | 9.0 | 9.0 | 9.0 | 9.0 |
| epochs | 33 | 50 | 21 | 108 |

## B Alignment

### B.1 Alignment Experiment

To perform alignment, we select a subset of 460 pocket pairs from the training set of the scoring neural network (section 2.3) and align them with RANSAC or our optimization-based approach described in section 2.2. For RANSAC, we consider three different modifications: RANSAC, RANSAC followed by point-to-point ICP (Besl & McKay, 1992), and RANSAC followed by point-to-plane ICP (Chen & Medioni, 1992). For our optimization-based method, we consider three objectives based on different underlying similarity functions: Euclidean distance, dot product, and cosine distance. To measure the quality of aligned pairs, we compute the root-mean-square deviation (RMSD) between ligand atoms that are transformed according to the corresponding pocket alignment. We report initial RMSD between atoms of ligands (i.e. before alignment), and final RMSD computed on transformed ligands in Table S8. We also provide the fraction of pairs for which alignment improved RMSD between atoms of ligands. For each method reported in Table S8, we chose the best set of hyperparameters. For details, see Appendix B.2. Figure S3 shows the RMSD distribution over all pairs before and after alignment. The distribution of errors on pockets aligned by RANSAC is shifted towards zero most, which makes this method the most preferable. On average, alignment of one pocket pair takes 0.17 seconds for RANSAC and 2.33 seconds for our optimization-based algorithm.

**Table S8:**
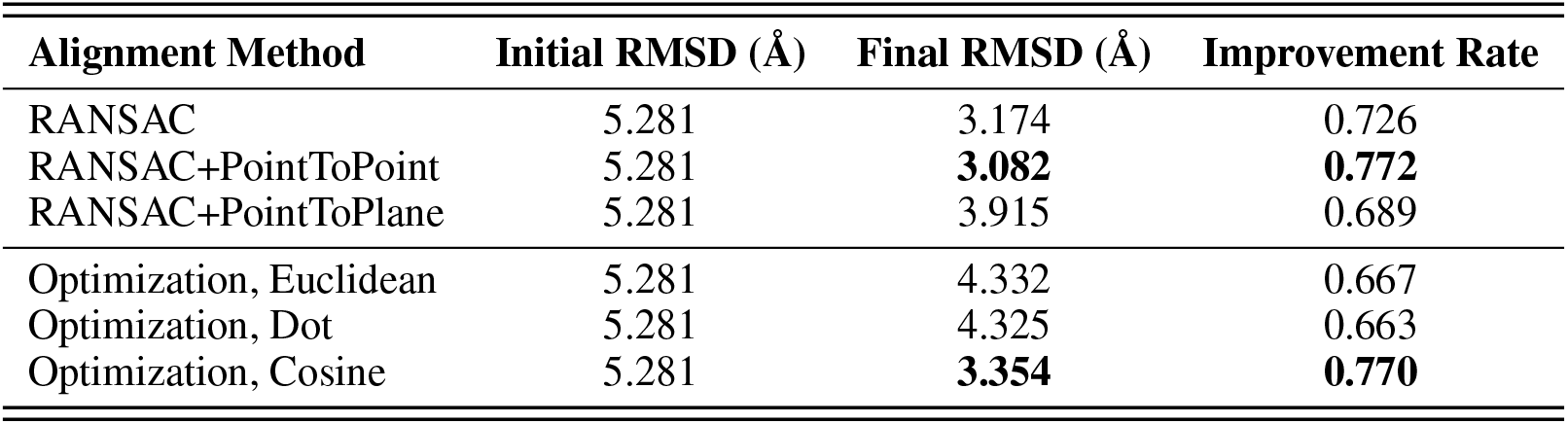
Alignment results.

| Alignment Method | Initial RMSD (Å) | Final RMSD (Å) | Improvement Rate |
| --- | --- | --- | --- |
| RANSAC | 5.281 | 3.174 | 0.726 |
| RANSAC+PointToPoint | 5.281 | **3.082** | **0.772** |
| RANSAC+PointToPlane | 5.281 | 3.915 | 0.689 |
| Optimization, Euclidean | 5.281 | 4.332 | 0.667 |
| Optimization, Dot | 5.281 | 4.325 | 0.663 |
| Optimization, Cosine | 5.281 | **3.354** | **0.770** |

**Figure S3:**
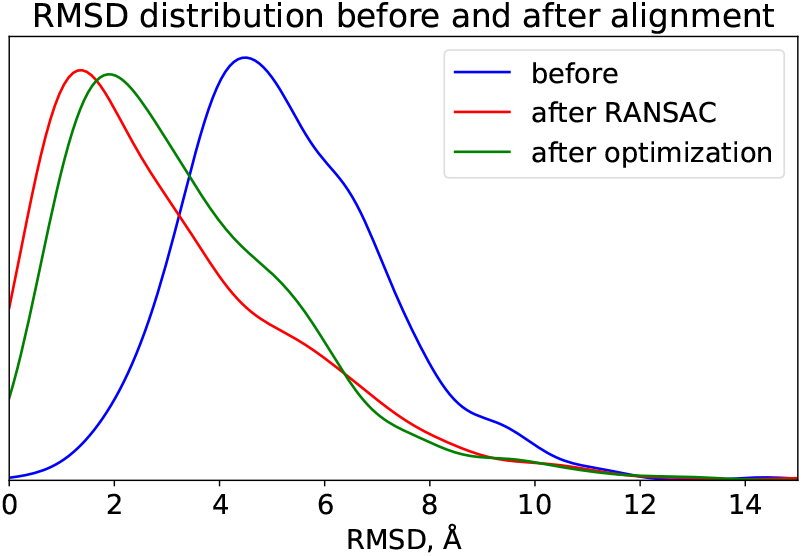
Distribution of RMSD values over pocket pairs before (blue) and after alignment (red and green).

### B.2 Alignment Parameters

In order to align pockets with RANSAC and ICP, we used the Open3D (Zhou et al., 2018) implementations of these algorithms. We considered different distance threshold values (between 1 Å and 10 Å) and different combinations of RANSAC and ICP: RANSAC, RANSAC+PointToPoint and RANSAC+PointToPlane. The final parameter set with the best performance is reported in Table S9.

**Table S9:** Best configuration of RASNAC+ICP parameters.

| ICP | distance_threshold | ransac_n | max_iter | max_validation |
| --- | --- | --- | --- | --- |
| PointToPoint | 2.0 | 3 | 100000 | 10000 |

For our optimization-based alignment algorithm, we considered three different similarity metrics (Euclidean distance, dot product, and cosine) and different values for the variance *σ* ∈ {0.5, 1.0, 2.0, 5.0, 10.0} of Gaussian kernels (1). We also varied *β* ∈ {0.0, 0.25, 0.5, 0.75, 1.0} in order to study the contribution of embeddings- and normals-related terms (2) to the total objective. Before alignment, we matched point clouds based on their centers of mass. We optimized translation parameter and rotation parameters with different learning rates, *lr*_*t*_ and *lr*_*R*_ correspondingly. Once the initial matching by centers of mass is performed, translation should not differ much from zero furthermore. Hence, the learning rate for the translation parameter was usually several orders of magnitude lower than learning rate for rotation parameters. In all experiments, we performed 1,000 steps of optimization. The best parameter configurations for each similarity function along with final RMSD results are provided in Table S10.

**Table S10:** Best parameter configurations of the optimization-based alignment algorithm with different similarity functions.

| Similarity | $\sigma$ | $\beta$ | $lr_t$ | $lr_R$ | Final RMSD ↑ |
| --- | --- | --- | --- | --- | --- |
| Euclidean | 10.0 | 0.75 | 0.0001 | 0.1 | 4.332 |
| Dot | 10.0 | 0.75 | 0.0001 | 0.1 | 4.325 |
| Cosine | 5.0 | 1.0 | 0.0001 | 1.0 | **3.354** |

## C Scoring Neural Network

The neural network was trained to solve the binary classification problem using cross-entropy loss. We train the network for 100 epochs with batch size 128 using Adam optimizer (Kingma & Ba, 2014) with learning rate 10^−4^.

## D Fragments

**Figure S4:**
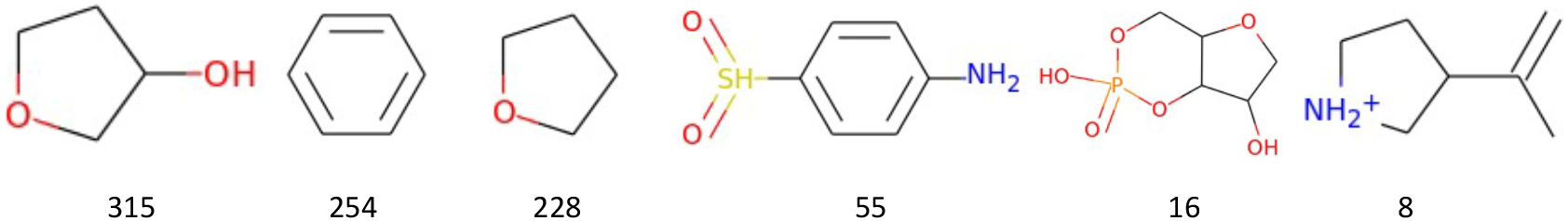
Examples of fragments retrieved using BRICS. Occurrences of fragments in our data are denoted beneath fragment representations.

## E Binding Affinity Prediction

### E.1 Hyperparameters

**Table S11:** Hyperparameters considered for binding affinity predictor.

| Component | Hyperparameter | Values |
| --- | --- | --- |
| **Surface Encoder** | embedding_layer | dMaSIF |
|  | resolution | 1.0 |
|  | distance | 1.05 |
|  | variance | 0.1 |
|  | sup_sampling | 10, **20** |
|  | atom_dims | 6 |
|  | emb_dims | 32 |
|  | in_channels | 16 |
|  | orientation_units | 16 |
|  | post_units | 8 |
|  | n_layers | **1**, 3 |
|  | radius | **9.0**, 15.0 |
| **Ligand Encoder** | Layers | [GCN, GCN, GCN, Linear] |
|  | Dims | [32, 32, 32, 32], [**128**, **128**, **128**, 32] |
| **Decoder** | Dims | [128, 128, 128, 1], [1024, 512, 128, 1], [**1024**, **1024**, **1024**, 1] |
|  | Batchsize | 128 |
|  | dropout | 0.3 |
|  | lr | 0.00001, **0.0001**, 0.001 |
| optimiser | Adam |  |

### E.2 Comparison on poses from co-crystallised structures

**Table S12:** Binding affinity prediction performance on PDBbind v2016 Core Set Docking Poses. [*] results taken from Jones et al. (2021). (R) and (G) denote models trained on the refined and general sets of PDBbind v2016.

| Method | RMSE ↓ | MAE ↓ | R ↑ |
| --- | --- | --- | --- |
| SG-CNN (R)* | 1.650 | 1.321 | 0.666 |
| SG-CNN (G)* | 1.508 | 1.277 | 0.699 |
| SG-CNN (R + G)* | 1.375 | 1.084 | 0.782 |
| 3D-CNN (R)* | 1.501 | 1.164 | 0.723 |
| 3D-CNN (G)* | 1.655 | 1.294 | 0.649 |
| 3D-CNN (R + G)* | 1.688 | 1.334 | 0.677 |
| Pafnucy* (Stepniewska-Dziubinska et al., 2018) | 1.42 | 1.13 | 0.78 |
| Kdeep* (Jiménez et al., 2018) | **1.27** | - | **0.82** |
| FAST-midlevel* (Jones et al., 2021) | 1.308 | **1.019** | 0.810 |
| FAST-late* (Jones et al., 2021) | 1.326 | 1.044 | 0.808 |
| Ours | 1.540 | 1.248 | 0.721 |

